# A Proposal for Finite But Unbounded Human Lifetimes

**DOI:** 10.1101/2022.01.07.474862

**Authors:** Fei Huang, Ross Maller, Brandon Milholland, Xu Ning

**Affiliations:** School of Risk & Actuarial Studies, UNSW Sydney; Research School of Finance, Actuarial Studies & Statistics, Australian National University; IQVIA, Plymouth Meeting, PA

## Abstract

Close analysis of an extensive data set combined with independent evidence prompts our proposal to view human lifetimes as individually finite but collectively unbounded. We formulate a model incorporating this idea whose predictions agree very well with the observed data. In the model, human lifetimes are theoretically unbounded, but the probability of an individual living to an extreme age is negligible, so lifetimes are effectively limited. Our model incorporates a mortality hazard rate plateau and a late-life mortality deceleration effect in conjunction with a newly observed advanced age mortality acceleration. This reconciles many previously observed effects. The model is temporally stable: consistent with observation, parameters do not change over time. As an application, assuming no major medical advances, we predict the emergence of many individuals living past 120, but due to accelerating mortality find it unlikely that any will subsequently survive to an age of 125.

## Introduction

The question of whether or not a limit to human lifespan exists may be as old as civilization itself. Earliest speculation was hampered by a lack of reliable age data, biological knowledge and statistical methods. Foundations for the modern field of demography were laid in the 19^*th*^ Century by Gompertz (*1*), who modelled mortality as increasing exponentially with age, and Makeham (*2*), who added an age-independent term to the equation. The Gompertz-Makeham Law of Mortality now enjoys near-universal acceptance as an accurate model for mortality at most adult ages. However, scientists still fiercely debate whether mortality deviates from the Gompertz-Makeham law at very old ages and whether human longevity can be increased into the future. On these issues, there is no consensus, with some foreseeing many years of extended longevity (*3*), while others argue that previous upward trends are largely played out and only modest improvements can be expected in the future (*4*).

In (*5*) an actuarially-motivated “smooth threshold life table” (STLT) model was fitted to a high-quality Netherlands data set: mortality data on 231,129 females and 73,788 males in 1-year, year-at-birth cohorts, from 1893 to 1908, with individuals aged up to 114 years (the recorded highest attained age). Fig. 4 in the Materials and Methods section displays the models fitted to the Netherlands data. The analysis was repeated on a smaller but up-to-date Australian data set reported in (*6*).

Herein the STLT model is closely analyzed to reveal fine structure details of the human mortality curve. An important feature found is an acceleration of mortality at very advanced ages. Based on this and other information from the modeling and the literature, we draw the plausible inference that individual human lifetimes are *inherently finite*.

Independent evidence in support of this can be found in an analysis of trends in three public longevity data sets (*7*), where it is argued that, nevertheless, *no hard upper bound* should be assumed for life-lengths overall. A continuing increase in human life expectancy since 1900 was deduced, but with a rate of improvement in survival which peaks and then declines for very old ages. It was inferred from this that human lifespan may have a natural limit but *maximum lifetime is not fixed* — although survival becomes negligible after a certain age, and there is no discernible improvement in extreme old-age survival over time.

Given this background, we argue that what is called for is a model in which life-lengths, though finite individually, vary randomly over the population according to a distribution which is unbounded. Herein we postulate such a model, estimate its parameters from the available data, and use it to predict numbers in the population who will live to very extreme ages. These predictions concur very well with the actual numbers observed.

## Results

### The STLT model

The STLT model in (*5*) and (*6*) consists of a Gompertz distribution describing lifelengths up till a data determined threshold age *N*, together with a generalised Pareto distribution (GPD) for ages greater than *N*. It incorporates a “late-life mortality deceleration effect”, where the rate of increase in mortality with advanced age becomes slower than exponential (as it is in the Gompertz law), and a “mortality hazard rate plateau” at high ages. Such effects have been observed before ((*8*), (*9*), (*10*)); see the Materials and Methods section for further discussion.

But how can a late-life mortality deceleration effect, for which mortality will tend to decrease and even level off at high ages, be consistent with finite human lifetimes? The answer is that the STLT model also implies an *advanced age mortality acceleration* following the late life mortality deceleration. This acceleration in hazards results in finite limits to the life spans. In the following we develop this idea.

### Finite but unlimited human lifetimes

The GPD formulation contains a shape parameter *κ* which is inversely proportional to the right extreme of the fitted mortality distribution. (See the Materials and Methods section for details.) For convenience, we take the shape parameter *κ* in the GPD to be the negative of that in the usual formulation (the *ξ* in (*11*), p.162, and the *γ* in (*12*) and (*13*)). A positive *κ* value signifies a finite upper bound to the lifetime distribution. The *κ* values estimated from the 32 year-of-birth *×* sex cohorts in the Netherlands data and for the 18 5-year Australian cohorts are all significantly greater than 0. They vary over a range shown in Table 1 for the Netherlands data and in Tables 4 and 5 of the Materials and Methods section for the Australian data.

**Table 1:** STLT parameter estimates and derived quantities. Û is the estimated upper bound age. All = all cohorts pooled.

| Cohort | Females |  |  |  |  | Males |  |  |  |  |
| --- | --- | --- | --- | --- | --- | --- | --- | --- | --- | --- |
| | $\hat{\kappa}$ | $\hat{N}$ | $x_A$ | $x_B$ | $\hat{U}$ | $\hat{\kappa}$ | $\hat{N}$ | $x_A$ | $x_B$ | $\hat{U}$ |
| 1893 | 0.159 | 97 | 105.91 | 109.34 | 114.74 | 0.221 | 89 | 100.62 | 104.46 | 109.55 |
| 1894 | 0.160 | 98 | 104.91 | 108.04 | 114.13 | 0.195 | 91 | 100.98 | 104.72 | 110.42 |
| 1895 | 0.205 | 98 | 100.64 | 102.48 | 110.67 | 0.226 | 89 | 99.99 | 103.76 | 109.00 |
| 1896 | 0.173 | 98 | 103.55 | 106.39 | 112.95 | 0.176 | 98 | 98.73 | 99.39 | 110.38 |
| 1897 | 0.207 | 97 | 101.30 | 103.77 | 110.88 | 0.179 | 98 | 98.49 | 98.95 | 110.37 |
| 1898 | 0.154 | 98 | 106.19 | 109.44 | 114.77 | 0.151 | 98 | 100.86 | 102.89 | 112.27 |
| 1899 | 0.171 | 98 | 104.10 | 107.01 | 113.10 | 0.179 | 95 | 99.57 | 102.31 | 110.82 |
| 1900 | 0.208 | 98 | 100.77 | 102.64 | 110.24 | 0.134 | 96 | 104.96 | 108.69 | 115.17 |
| 1901 | 0.191 | 97 | 103.06 | 105.91 | 111.78 | 0.181 | 94 | 100.08 | 103.28 | 111.12 |
| 1902 | 0.151 | 98 | 106.76 | 109.98 | 114.78 | 0.109 | 97 | 108.98 | 113.04 | 118.56 |
| 1903 | 0.133 | 98 | 109.38 | 112.70 | 116.59 | 0.116 | 98 | 106.49 | 110.19 | 117.01 |
| 1904 | 0.174 | 96 | 106.13 | 109.37 | 113.52 | 0.197 | 93 | 98.69 | 101.78 | 109.65 |
| 1905 | 0.144 | 97 | 109.19 | 112.48 | 116.03 | 0.139 | 98 | 102.49 | 105.22 | 113.87 |
| 1906 | 0.135 | 98 | 109.04 | 112.26 | 116.02 | 0.125 | 95 | 107.95 | 112.07 | 117.31 |
| 1907 | 0.166 | 97 | 105.93 | 109.02 | 113.35 | 0.145 | 98 | 101.51 | 103.83 | 112.92 |
| 1908 | 0.208 | 95 | 103.74 | 106.80 | 111.11 | 0.134 | 96 | 104.76 | 108.45 | 114.94 |
| All | 0.151 | 98 | 106.52 | 109.75 | 114.74 | 0.151 | 97 | 101.49 | 104.19 | 112.69 |

We infer that lifetimes in a cohort can be modelled as bounded above at a finite age *U* inversely proportional to *κ*, but with *U* varying randomly over the cohorts. We extend this insight to a model in which individual lifetimes are finite but with upper bounds which vary over the population with no definite upper limit.

### A mixture model for lifetimes

Write the lifetime *T* of a person in the population as *T* = *N* + *Y*, where *Y* is the excess lifetime above the transition age *N*. Thus *Y* has as distribution the Pareto distribution component of the STLT model, conditional on having reached age *N*, with shape parameter the *κ* in that component.

Assume that associated with each person is a positive random variable whose realized value is *κ*. Then the lifetime distribution in the population as a whole is given by integrating the conditional distribution over possible values of *κ*. The distribution of *κ* required for this can be estimated from the fitted STLT models. We used the *κ* values in Table 1 (Netherlands data) combined with those in Tables 4 and 5 in the Materials and Methods section (Australian data). The resulting empirical distribution is shown in Fig. 5 of the Materials and Methods section. We approximated the histogram in Fig. 5 with a gamma(*a, b*) distribution. The resulting distribution of *Y* calculated by mixing over the distribution of *κ* using the estimated density for *κ* is in Eq. (8) of the Materials and Methods section. It is unbounded, consistent with our belief that there is no hard upper bound on lifetimes overall. The resulting population distribution densities are shown in Fig. 1. There are non-negligible probabilities of values of *Y* up till about 12, corresponding to *T ≈* 98 + 12 = 110 years. Survival probabilities then decline rapidly and become negligible by age 120.

**Figure 1:**
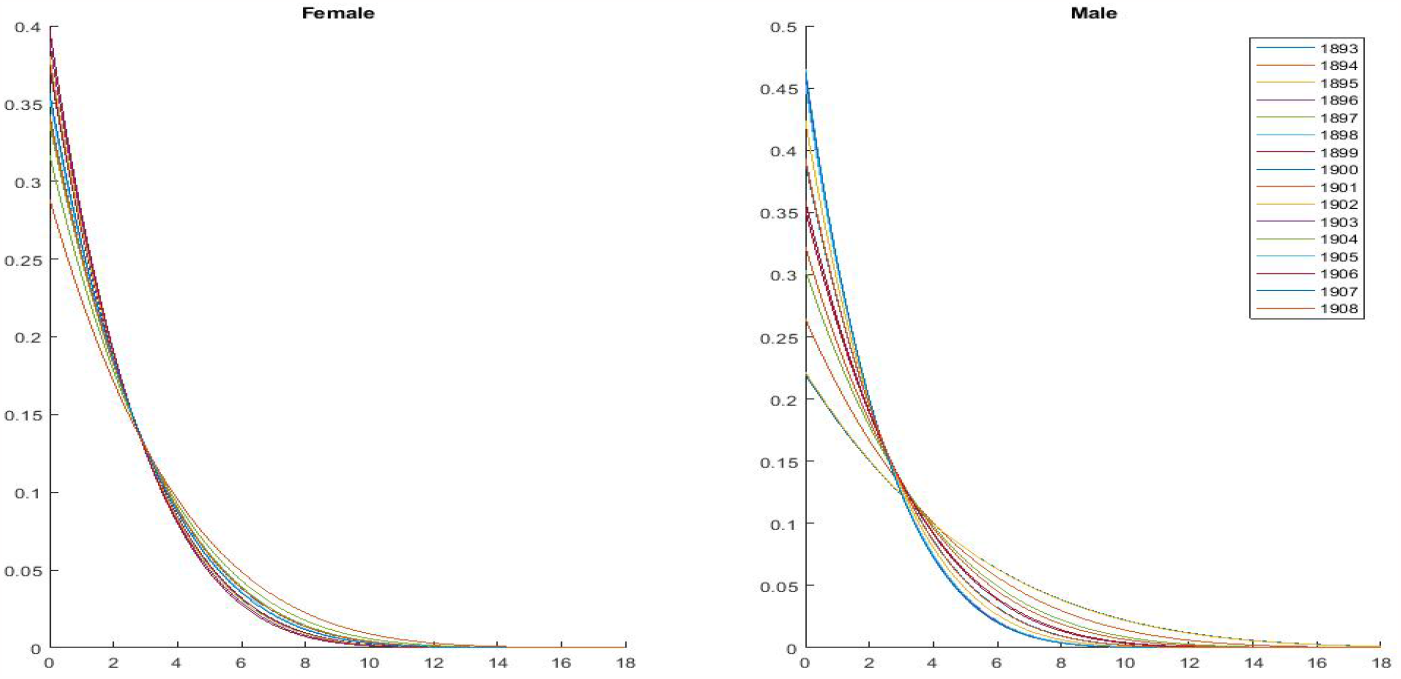
Lifetime densities above transition age for 32 Netherlands cohorts.

### Advanced age mortality accelerations

The existence of an age *x*_*A*_ before which the force of mortality (the hazard) in the GPD increases slower than that of the Gompertz, but after which the GPD hazard increases faster than that of the Gompertz, is verified for the STLT model in the Materials and Methods section. This phenomenon gives rise to the advanced age mortality acceleration. In the Netherlands data, the acceleration is observed subsequent to the threshold age *N* and prior to the limiting age *U* for all cohorts. The ages *x*_*A*_ are listed for each cohort in Table 1 for pooled data (all Netherlands cohorts combined, separately for females and males). Figure 2 shows Kaplan-Meier estimates (KMEs) for survival curves calculated from this data together with the fitted STLT models (left panel) and the corresponding hazard functions and hazard function slopes derived from the fitted curves (right hand panels). The baseline values for the KMEs in the left panel are the empirical values of the actuarial mortality estimates at age 65 for Netherlands females and males (all cohorts combined). The fitted models provide an excellent description of the data; for more detail on this see Fig. 4 in the Materials and Methods section. The ages *x*_*B*_ at which the hazards of the Gompertz and Pareto curves cross again are also shown in Table 1 and Figures 2 and 4. Further information on the advanced age mortality acceleration and its relation to the “three laws of biodemography” is in the Discussion

**Figure 2:**
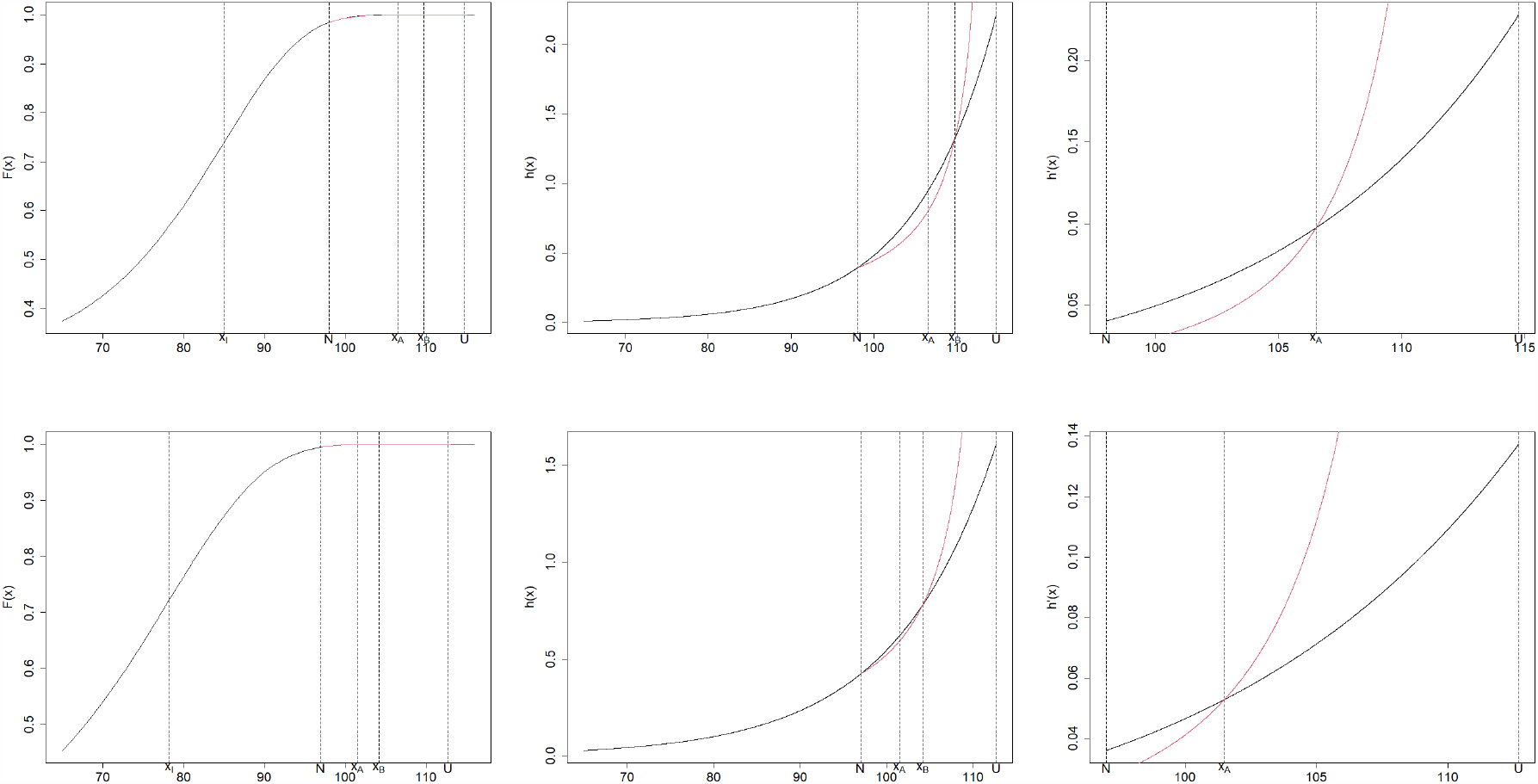
Upper panel: female; lower panel: male. Left: observed and fitted cdfs with mortality plateau ≈ age 105+; Middle: corresponding hazard functions; Right: hazard function slopes, showing late life mortality deceleration for ages N < x < x_A_ and advanced age mortality acceleration for ages x > x_A_.

### Observed vs expected centenarians

The probability of living to age *x > N* years or more can be calculated as

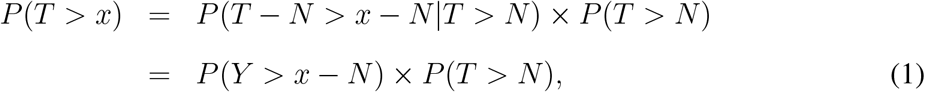

where *N* is the transition age for the cohort under consideration. Multiplying the probability *P* (*Y > x − N*) in Eq. (1) by the number in the cohort at age *N* gives the expected number in the cohort living to age *x*, or beyond, according to the model. These are compared with the actual numbers for ages *x* = 100, 105 in Fig. 3. (The numbers themselves, also for *x* = 110, are in Table 6 of the Materials and Methods section. No person of either sex in the data lived longer than 115 years, and at this age model predictions were *<* 0.01 persons.)

**Figure 3:**
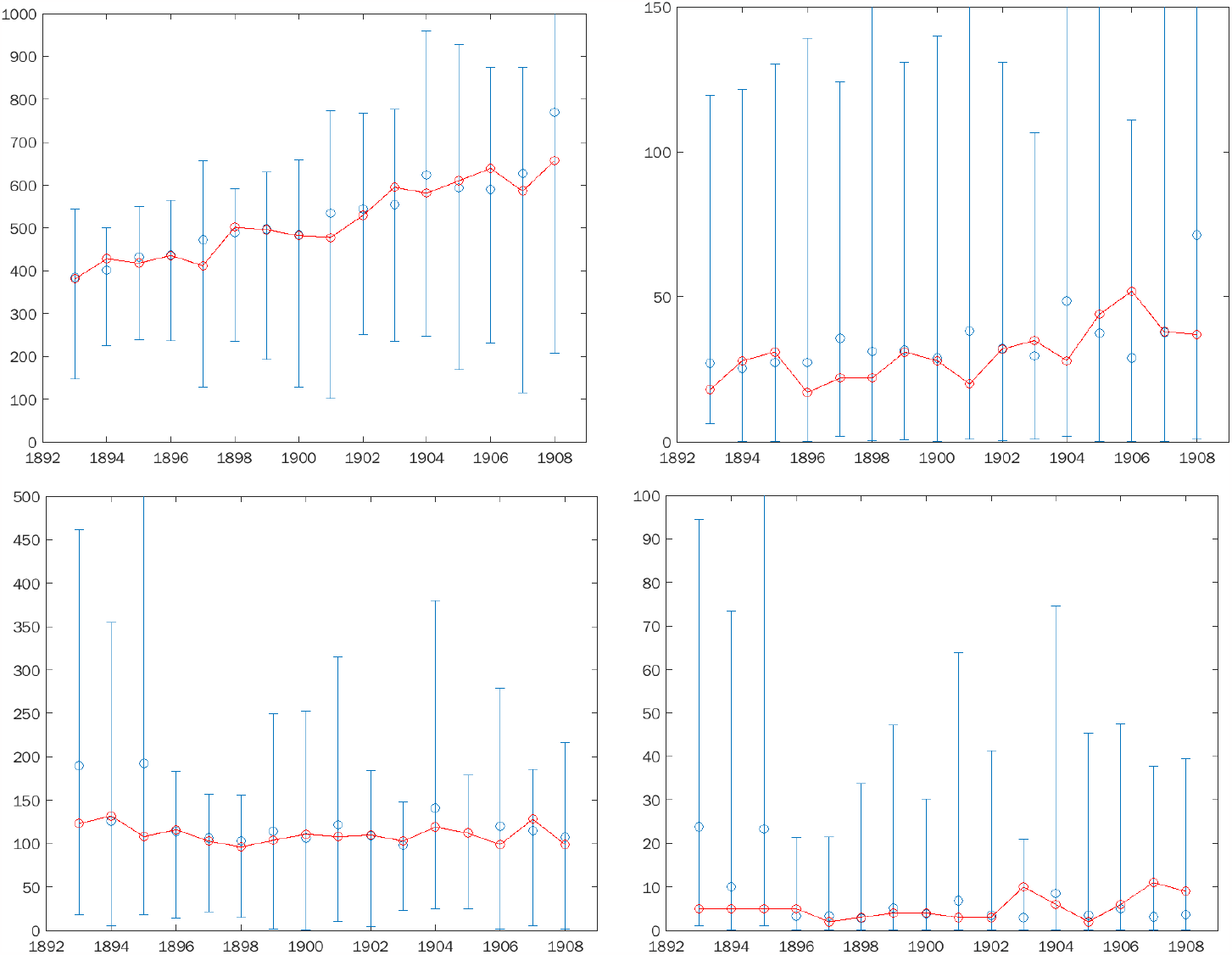
Numbers of observed (red) and predicted by model (blue) living to ages ≥ 100 (left column) and ≥ 105 (right column) in cohorts 1893–1906. Upper panel: female; lower panel: male. Approximate 95% confidence intervals around the estimated numbers were calculated by resampling.

**Figure 4:**
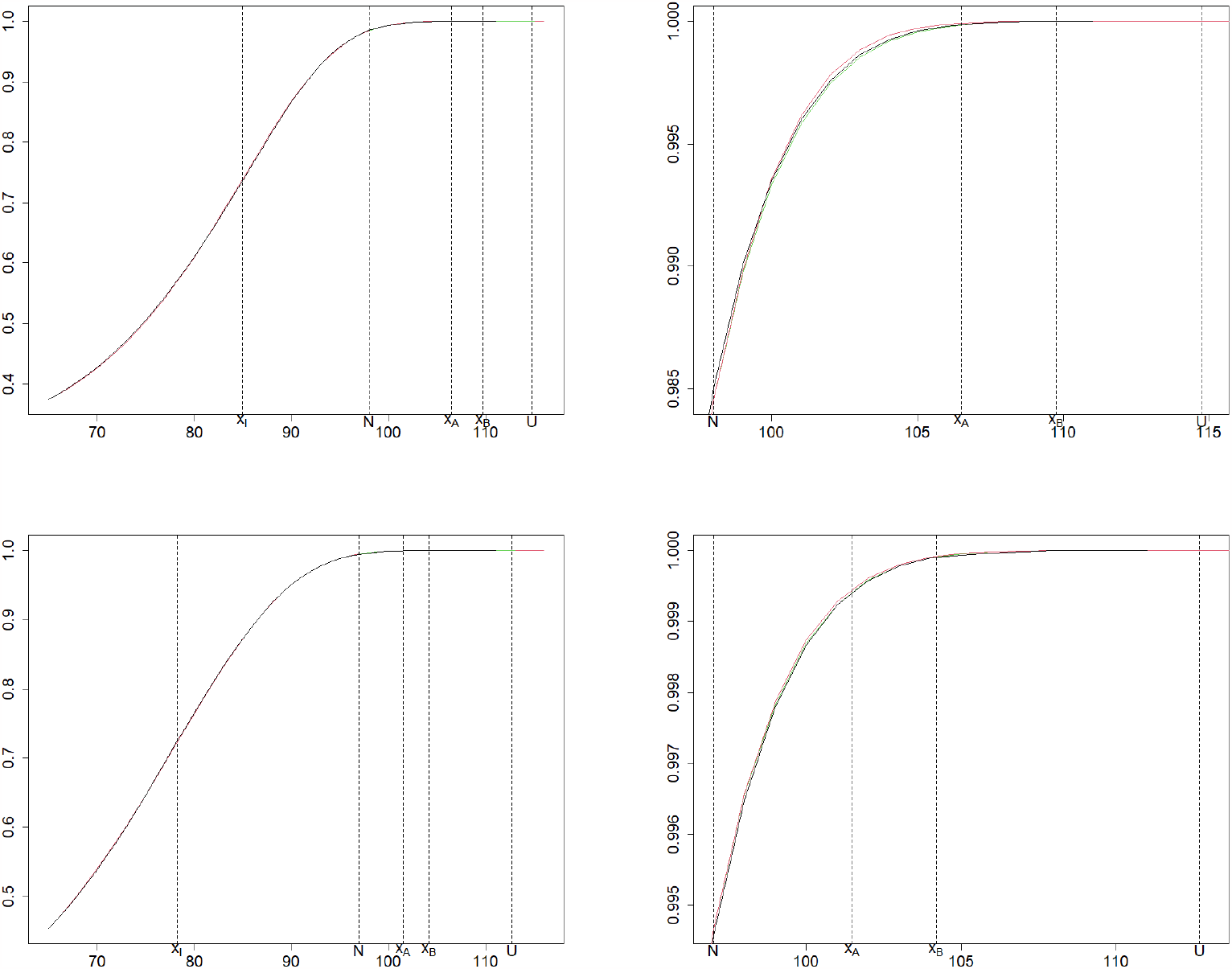
Black lines: KMEs for Netherlands data, all cohorts. Upper panel: female; lower panel: male. Left column, all ages 65+, with red: fitted Gompertz, ages < N, and green: fitted Pareto, ages ≥ N. Right column, ages ≥ N only; red is Gompertz extrapolated to ages ≥ N.

**Figure 5:**
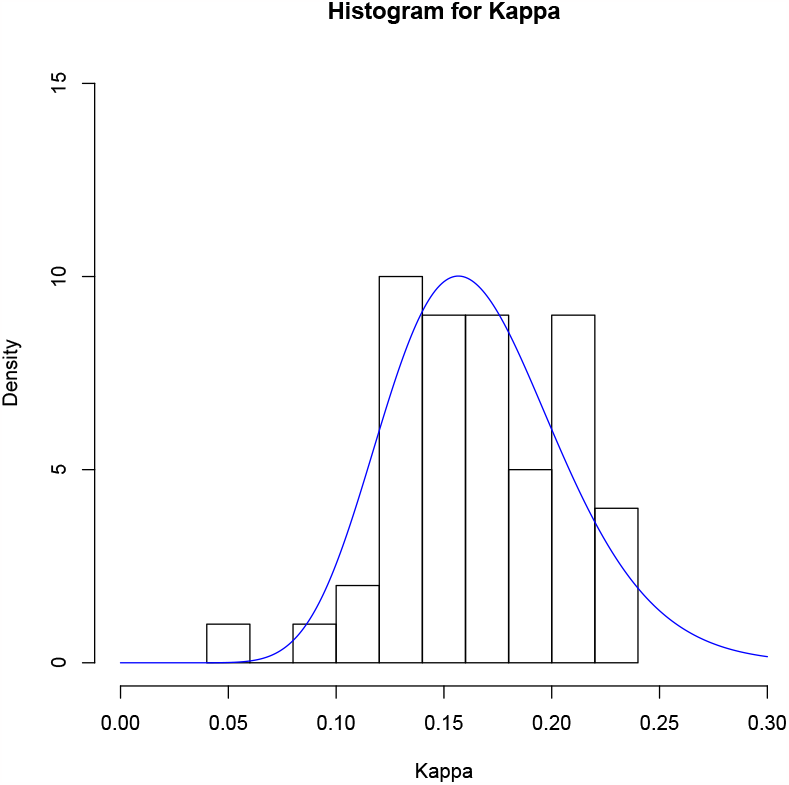
Histogram of combined Netherlands and Australian κ values with fitted gamma(a, b) density. Maximum likelihood estimates and SEs are â = 16.63 (0.81) and 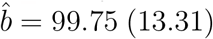.

Observed and expected compare well but some inaccuracy is to be expected especially for the smaller numbers as we are attempting to predict in the extreme upper end of the life-table. We note an increasing trend with birth cohort year for females surviving to 100 years or beyond, but not for the other groups.

### Expected survivors at extreme ages

How many individuals exceeding Jeanne Calment’s record age (122 years and 164 days) might we expect to see in the remainder of this century? In the mixture model, while survival to age 115 or older is possible, it is with a probability so small as to be negligible. One could speculate that, on a global scale, an increasing number of supercentenarians might result in the appearance of new longevity records. However, our findings place a robust limit on the emergence of any record-breaking individuals.

Assuming the model derived from the Netherlands and Australian data applies generally, we can estimate roughly the potential growth in the number of supercentenarians, globally, from 2021-2100. The annual growth in supercentenarians recorded in the International Database on Longevity (IDL) for England and Wales, 1968–2017, is well described by an exponential growth function with a doubling-time of 12.9 years (see the Materials and Methods section). From this function we extrapolated, optimistically, the supercentenarian count to be expected in England and Wales for 2021–2100. Multiplying by a factor of 100 (for the approximate proportion of the world population in England and Wales) gives an estimate of 1,476,796, or in broad terms a total of *≈* 1.5 million supercentenarians, male and female, to be expected world-wide over the 80-year period 2021–2100.

For an optimistic estimate of the number of survivors from this population expected to exceed a specified age of interest, we multiply 1.5*×*10^6^ by the corresponding proportion surviving calculated from the (female) density values in Fig. 1. The model then predicts that 8268 of those 1.5m supercentenarians would survive to age 115, and 41 would live to age 120, but less than 1 (expectation 0.23) would live past age 125. These rough estimates at the extreme upper end of the age distribution come with large standard errors, but are consistent with the small numbers of extreme-aged individuals currently observed, and suggest there is a low probability of Jeanne Calment’s record being broken even once in the remainder of this century. Barring a revolutionary advance in medicine, the benchmark age of 125 appears to represent a firm limit to human longevity unlikely to be exceeded in the foreseeable future.

## Discussion

### Is there an upper limit to human lifetimes?

Much commentary on this issue related to (*7*), some critical, some supportive, followed soon after its publication. A study based on the IDL alone (*13*) was critical: it rejected the inferred finite limit to human life, and, using longevity data from Japan and Western countries, suggested instead an exponential model for lifetimes beyond 110 years. As in (*5*), the study fitted the GPD, but in this case, to data from 4 areas (North and South Europe, North America and Japan), and to a very limited set (a total of 566 life lengths) from extreme ages only (supercentenarians, 110+). In two instances (North Europe and Japan) the estimated *κ* in their GPD fits is negative (we write *κ* for *−γ*, in their notation), while for South Europe and North America it is positive. But in no case is it significantly different from 0 (Table 6, (*13*)) and pooled *κ* estimates are all positive. From being non-significantly different to 0, the authors concluded that *κ* should be taken as exactly zero, indicating an exponential distribution for the data, and argue from this for an unlimited human lifespan.

Later (*14*) and (*15*) updated (*13*) with more data and analyses but ultimately still concluded in favour of an exponential model for lifetimes and an unlimited human lifespan.

However, these analyses obtaining non-significant positive (and in some cases non-significant negative) *κ* values are in fact consistent with those in (*5*) and (*6*), where the estimated *κ* are strongly positive. Only limited data from extreme ages is used in (*13*), (*14*) and (*15*), whereas in (*5*) and (*6*) more extensive data sets spanning ages from 65 on was used, thus obtaining greater statistical power in a demographically-informed model.

Other comments on (*7*) occurred in (*16*), (*17*), (*18*), and (*19*). Most of those writers argued for the contrary conclusion – for no upper limit to human lifespan. One comment (*20*), using a logistic model, argued that an individual may live to 125 years within this century. Their model tacitly assumes an infinite endpoint.

By contrast, a number of papers which also include discussion or estimation of the GPD shape parameter *κ* come to similar conclusions as (*7*), (*5*), and (*6*). They include: an analysis of data from the Gerontology Research Group, which also observed no trend in supercentenarian longevity (*21*); an actuarial study of 46,66 Belgians *>* 95 years old, finding a limit to lifetimes of *≈* 120 *−* 130 years and no trend across cohorts (*22*); an analysis of the combined datasets of the IDL and the Human Mortality Database, which found a limit of *≈* 125 *−* 130 years (*23*); and several reanalyses of the IDL in response to (*13*): (*24*), (*25*), (*26*), and (*27*). Finally, a study of *≈* 285,000 Netherlands individuals aged 92 years or more at death consistently found positive estimates of *κ*, as well as a limit of *≈* 115 *−* 130 years for maximum human lifespan, with no trend over time (*12*).

Almost all papers, even those which estimated a negative *κ* value, find that the probability of survival becomes either exactly 0 or negligibly higher than 0 at some point between 115–130 years. Indeed, the classification of lifespan as “unlimited” based on a non-zero yet extremely small chance of survival has been criticized previously (*28*). Our analyses confirm and strengthen this consensus and refute other analyses that predict longevity exceeding 125 years (*20*). Even assuming unabated exponential growth in the number of supercentenarians and applying an optimistic model of supercentenarian survival, we predict that the probability of observing an individual living past 125 by 2100 is 0.23.

But exponential growth in the number of supercentenarians past 2100 is improbable, anyway. The present exponential growth in supercentenarians is driven by two factors: growth in birth cohort sizes and increased survival to age 110. Birth cohort sizes peaked in the mid-20^*th*^ century for many nations and are unlikely to contribute to higher supercentenarian numbers during the 22^*nd*^ century. Meanwhile, the stagnation, or even decline, in life expectancy for developed nations indicates that the proportion of each birth cohort surviving to age 110 is unlikely to increase beyond this century. Together, these factors suggest that supercentenarian cohort sizes — and longevity records — will stop growing or even fall in the following century, if not sooner.

Alternatively, a breakthrough in medicine could result in greatly increased longevity at extreme ages. However, consideration of this is beyond the scope of our present analysis.

### Late-life mortality plateau and deceleration effect

An earlier study (*8*) tracked every person born in Italy between 1896 and 1910 who lived to age 105 or beyond; a total of 3,836 individuals (3,373 women and 463 men). That study’s hazard rate analysis suggested a “mortality hazard rate plateau” between ages 105 and 110 and a consequent “late-life mortality deceleration effect” – that is, that the rate of increase in mortality with advanced age becomes slower than exponential (as it is in the Gompertz law) between those ages. This effect is usually understood to imply a model with a potentially infinite life span. The existence of a late-life mortality hazard rate plateau is also argued by earlier studies (*10*), (*9*), but disputed and critiqued by several commentaries (*28*), (*29*), (*30*), (*31*). As we observed in the main text, the existence of an advanced age mortality acceleration resolves any contradiction of these studies with our conclusion of finite lifetimes.

### The three laws of biodemography

The compensation law of mortality ((*32*), (*33*), (*34*)) states that for a given species, differences in death rates between different sub-populations (cohorts, in our case) decrease with age — because higher or lower initial death rates are accompanied by a lower or higher rate of mortality increase with age, such that mortality rates for different cohorts tend to equalise after some high age.

This tendency exists in the Netherlands data and is captured in the STLT model fit. The threshold ages *N* at which the convergence takes place are around 98 years for females and (with somewhat greater variability) around 96 years for males (Tables 2 and 3). These are close to the estimate of around 95 years made in (*33*) on quite different grounds.

**Table 2:**
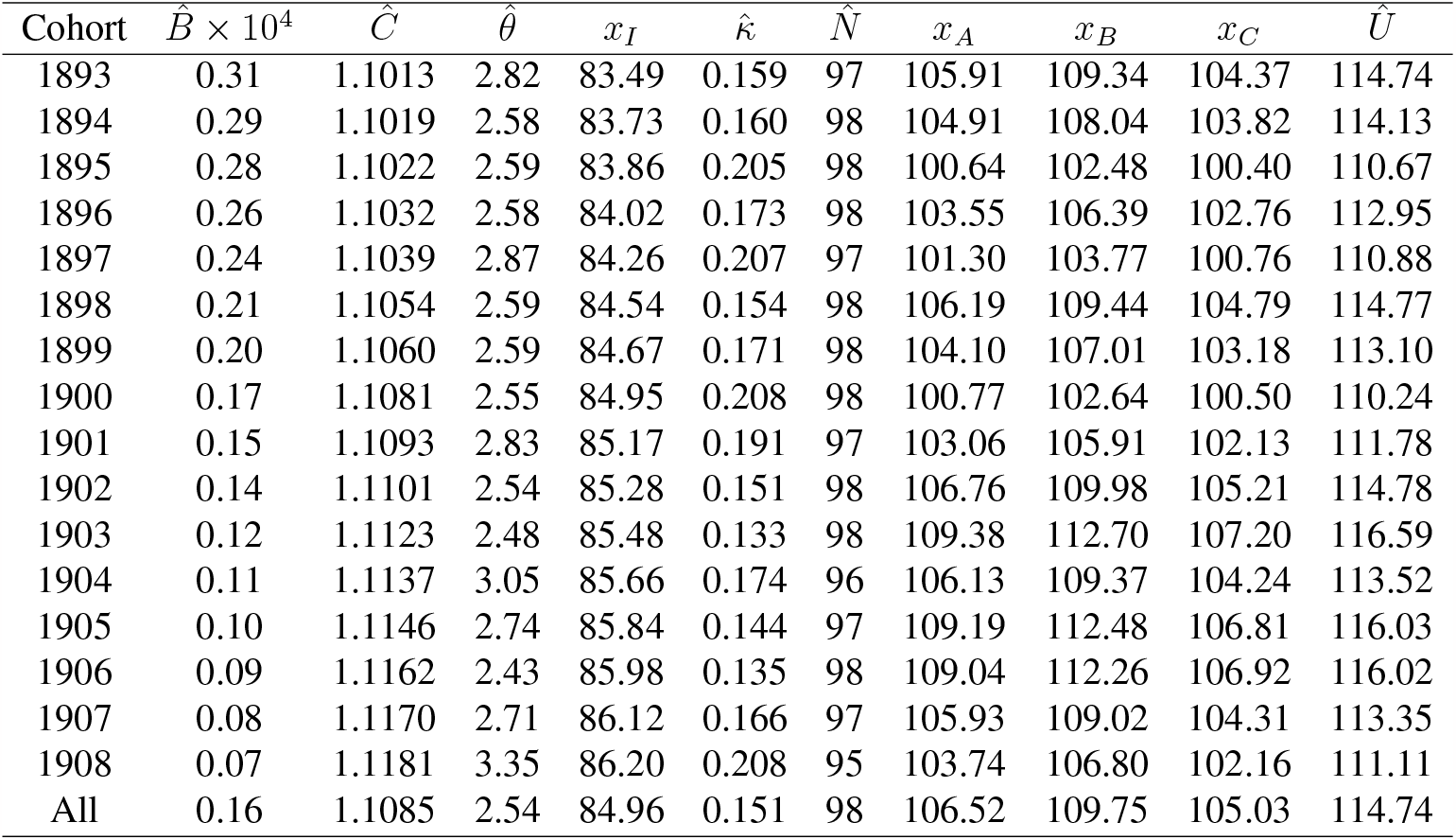
Estimated parameters for each birth cohort of Netherlands females, with estimated ages 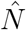, x_I_, x_A_, x_B_, x_C_, Û. All = data for all female cohorts pooled. Standard errors for the parameter estimates are in (5).

| Cohort | $\hat{B} \times 10^4$ | $\hat{C}$ | $\hat{\theta}$ | $x_I$ | $\hat{\kappa}$ | $\hat{N}$ | $x_A$ | $x_B$ | $x_C$ | $\hat{U}$ |
| --- | --- | --- | --- | --- | --- | --- | --- | --- | --- | --- |
| 1893 | 0.31 | 1.1013 | 2.82 | 83.49 | 0.159 | 97 | 105.91 | 109.34 | 104.37 | 114.74 |
| 1894 | 0.29 | 1.1019 | 2.58 | 83.73 | 0.160 | 98 | 104.91 | 108.04 | 103.82 | 114.13 |
| 1895 | 0.28 | 1.1022 | 2.59 | 83.86 | 0.205 | 98 | 100.64 | 102.48 | 100.40 | 110.67 |
| 1896 | 0.26 | 1.1032 | 2.58 | 84.02 | 0.173 | 98 | 103.55 | 106.39 | 102.76 | 112.95 |
| 1897 | 0.24 | 1.1039 | 2.87 | 84.26 | 0.207 | 97 | 101.30 | 103.77 | 100.76 | 110.88 |
| 1898 | 0.21 | 1.1054 | 2.59 | 84.54 | 0.154 | 98 | 106.19 | 109.44 | 104.79 | 114.77 |
| 1899 | 0.20 | 1.1060 | 2.59 | 84.67 | 0.171 | 98 | 104.10 | 107.01 | 103.18 | 113.10 |
| 1900 | 0.17 | 1.1081 | 2.55 | 84.95 | 0.208 | 98 | 100.77 | 102.64 | 100.50 | 110.24 |
| 1901 | 0.15 | 1.1093 | 2.83 | 85.17 | 0.191 | 97 | 103.06 | 105.91 | 102.13 | 111.78 |
| 1902 | 0.14 | 1.1101 | 2.54 | 85.28 | 0.151 | 98 | 106.76 | 109.98 | 105.21 | 114.78 |
| 1903 | 0.12 | 1.1123 | 2.48 | 85.48 | 0.133 | 98 | 109.38 | 112.70 | 107.20 | 116.59 |
| 1904 | 0.11 | 1.1137 | 3.05 | 85.66 | 0.174 | 96 | 106.13 | 109.37 | 104.24 | 113.52 |
| 1905 | 0.10 | 1.1146 | 2.74 | 85.84 | 0.144 | 97 | 109.19 | 112.48 | 106.81 | 116.03 |
| 1906 | 0.09 | 1.1162 | 2.43 | 85.98 | 0.135 | 98 | 109.04 | 112.26 | 106.92 | 116.02 |
| 1907 | 0.08 | 1.1170 | 2.71 | 86.12 | 0.166 | 97 | 105.93 | 109.02 | 104.31 | 113.35 |
| 1908 | 0.07 | 1.1181 | 3.35 | 86.20 | 0.208 | 95 | 103.74 | 106.80 | 102.16 | 111.11 |
| All | 0.16 | 1.1085 | 2.54 | 84.96 | 0.151 | 98 | 106.52 | 109.75 | 105.03 | 114.74 |

**Table 3:**
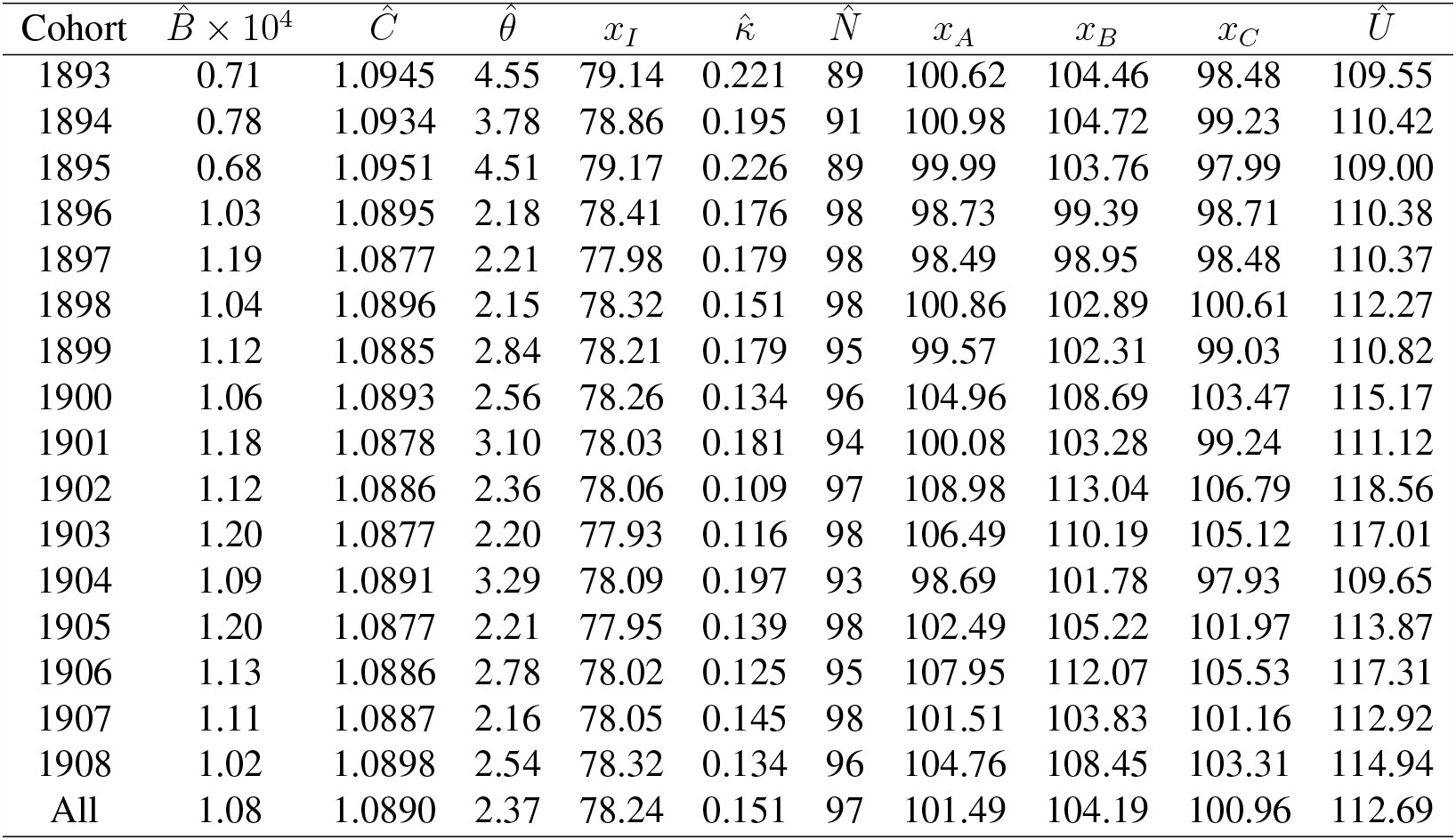
Estimated parameters for each birth cohort of Netherlands males, with estimated ages 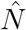, x_I_, x_A_, x_B_, x_C_, Û.All = data for all male cohorts pooled. Standard errors for the parameter estimates are in (5).

| Cohort | $\hat{B} \times 10^4$ | $\hat{C}$ | $\hat{\theta}$ | $x_I$ | $\hat{\kappa}$ | $\hat{N}$ | $x_A$ | $x_B$ | $x_C$ | $\hat{U}$ |
| --- | --- | --- | --- | --- | --- | --- | --- | --- | --- | --- |
| 1893 | 0.71 | 1.0945 | 4.55 | 79.14 | 0.221 | 89 | 100.62 | 104.46 | 98.48 | 109.55 |
| 1894 | 0.78 | 1.0934 | 3.78 | 78.86 | 0.195 | 91 | 100.98 | 104.72 | 99.23 | 110.42 |
| 1895 | 0.68 | 1.0951 | 4.51 | 79.17 | 0.226 | 89 | 99.99 | 103.76 | 97.99 | 109.00 |
| 1896 | 1.03 | 1.0895 | 2.18 | 78.41 | 0.176 | 98 | 98.73 | 99.39 | 98.71 | 110.38 |
| 1897 | 1.19 | 1.0877 | 2.21 | 77.98 | 0.179 | 98 | 98.49 | 98.95 | 98.48 | 110.37 |
| 1898 | 1.04 | 1.0896 | 2.15 | 78.32 | 0.151 | 98 | 100.86 | 102.89 | 100.61 | 112.27 |
| 1899 | 1.12 | 1.0885 | 2.84 | 78.21 | 0.179 | 95 | 99.57 | 102.31 | 99.03 | 110.82 |
| 1900 | 1.06 | 1.0893 | 2.56 | 78.26 | 0.134 | 96 | 104.96 | 108.69 | 103.47 | 115.17 |
| 1901 | 1.18 | 1.0878 | 3.10 | 78.03 | 0.181 | 94 | 100.08 | 103.28 | 99.24 | 111.12 |
| 1902 | 1.12 | 1.0886 | 2.36 | 78.06 | 0.109 | 97 | 108.98 | 113.04 | 106.79 | 118.56 |
| 1903 | 1.20 | 1.0877 | 2.20 | 77.93 | 0.116 | 98 | 106.49 | 110.19 | 105.12 | 117.01 |
| 1904 | 1.09 | 1.0891 | 3.29 | 78.09 | 0.197 | 93 | 98.69 | 101.78 | 97.93 | 109.65 |
| 1905 | 1.20 | 1.0877 | 2.21 | 77.95 | 0.139 | 98 | 102.49 | 105.22 | 101.97 | 113.87 |
| 1906 | 1.13 | 1.0886 | 2.78 | 78.02 | 0.125 | 95 | 107.95 | 112.07 | 105.53 | 117.31 |
| 1907 | 1.11 | 1.0887 | 2.16 | 78.05 | 0.145 | 98 | 101.51 | 103.83 | 101.16 | 112.92 |
| 1908 | 1.02 | 1.0898 | 2.54 | 78.32 | 0.134 | 96 | 104.76 | 108.45 | 103.31 | 114.94 |
| All | 1.08 | 1.0890 | 2.37 | 78.24 | 0.151 | 97 | 101.49 | 104.19 | 100.96 | 112.69 |

**Table 4:** Australian female cohort data.

| Year | $\hat{\kappa}$ | SE($\hat{\kappa}$) | $\hat{U}$ | SE($\hat{U}$) | $N$ |
| --- | --- | --- | --- | --- | --- |
| 1860 | 0.0591 | 0.0154 | 138.3107 | 11.2281 | 95 |
| 1865 | 0.2175 | 0.0040 | 109.9093 | 0.3497 | 88 |
| 1870 | 0.2061 | 0.0064 | 110.4127 | 0.5491 | 91 |
| 1875 | 0.0939 | 0.0138 | 125.6395 | 4.3085 | 96 |
| 1880 | 0.1220 | 0.0091 | 121.3737 | 1.9759 | 94 |
| 1885 | 0.2351 | 0.0043 | 111.1121 | 0.3405 | 89 |
| 1890 | 0.2043 | 0.0060 | 113.3055 | 0.5439 | 93 |
| 1895 | 0.2232 | 0.0058 | 112.2684 | 0.4304 | 94 |
| 1900 | 0.1281 | 0.0346 | 118.8452 | 4.2573 | 103 |

**Table 5:**
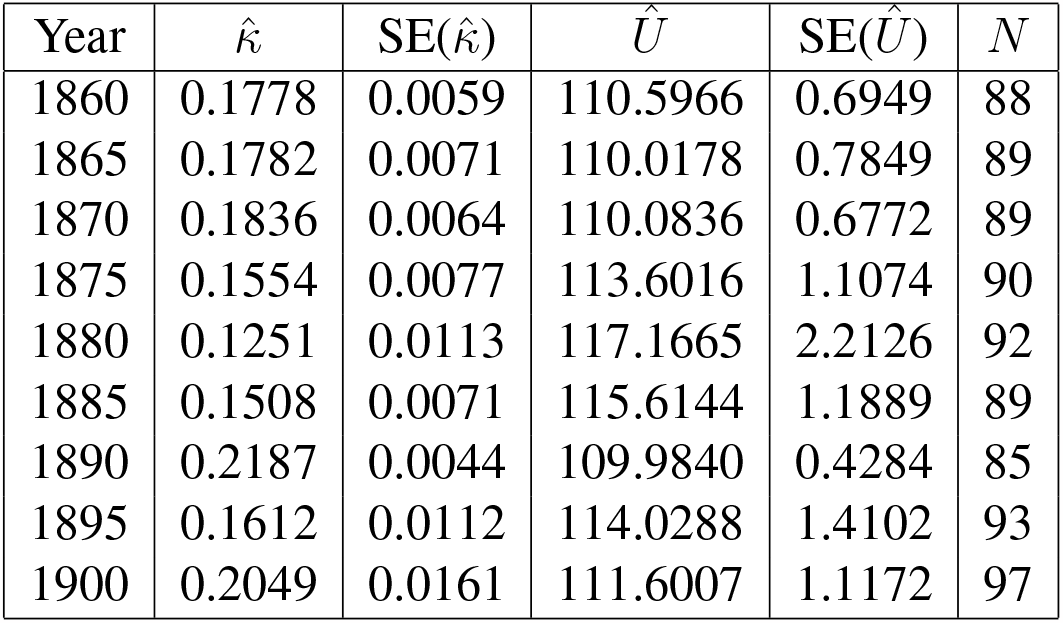
Australian male cohort data.

The three *laws of biodemography* have been stated as:

- the Gompertz-Makeham law of mortality;
- the compensation law of mortality, and
- the existence of a late-life mortality deceleration.

The Netherlands data and the STLT model fitted to it are consistent with the compensation law, so all three laws are satisfied. The model also allows for, and for that data, predicts, via an advanced age mortality acceleration, a finite limit to human life span. The STLT model is unique in possessing all of these properties.

The STLT model and the mixture model derived from it herein appear to represent the most biologically and statistically rigorous evaluation of late-life human mortality to date.

## Materials and Methods

### The STLT model: Gompertz & Pareto components

The model fitted in (*5*) and (*6*) to each of the cohorts in the Netherlands and Australian data consists of a Gompertz distribution with cumulative distribution function (cdf)

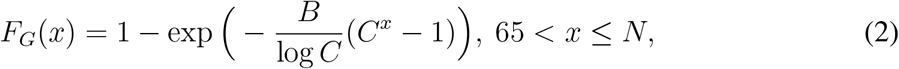

for parameters *B >* 0 and *C >* 1, describing a lifetime of length *x* up till a data determined threshold age *N*, together with a generalised Pareto distribution for ages greater than *N*. This Pareto part has the form ((*11*), p.162)

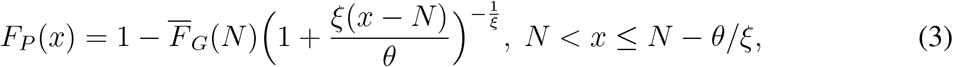

for parameters *θ >* 0 and *ξ*. (We denote the survival function (tail function) of a cdf *F* by 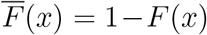, *x ≥* 0.) In general the parameter *ξ* can take either sign, but in the Netherlands data in (*5*) it was estimated to be significantly *negative* in all 32 cohorts, hence resulting in the finite upper bound for *x* indicated in Eq. (3). To avoid many minus signs in formulae, we write *κ* for *|ξ|* = *−ξ* throughout. With this notation the estimates of *κ* together with estimates of other significant parameters are in Tables 2 and 3 (Netherlands data) and Tables 4 and 5 (Australian data).

An innovative feature of the STLT model is that a smooth transition in hazard rates between ages less than and greater than the threshold age *N* is enforced. This means that the parameters in Eq. (2) and Eq. (3) are connected by the relation *θ* = 1*/BC^N^*.

We use the Kaplan-Meier empirical distribution estimator (KME) to display the lifetime data. Figure 4 shows the Netherlands data with the fitted STLT models (Gompertz up till the transition age *N*, Pareto thereafter), for females (upper panel) and males (lower panel). The left column is for all ages, 65+. The right column is a magnified view of the left column for ages greater than *N*. In this column the Gompertz distribution fitted to ages less than *N* is extrapolated to ages greater than *N*. Note that the extrapolated Gompertz tends to overestimate mortality slightly by comparison with the Pareto in this age range. The parameter estimates used to construct the curves are in Tables 2 and 3.

### STLT model: hazards

Formulae for the hazards of the Gompertz and Pareto components can be calculated from Eq. (2) and Eq. (3) and it can be checked that the STLT constraint forces *θ* = 1*/BC^N^* as claimed. Further analysis shows that there is a point *x*_*A*_ in (*N, U*) at which the hazards cross. For *x ∈* [*N, x_A_*) the Pareto hazard increases more slowly than the Gompertz hazard; but for *x ∈* (*x*_*A*_, *U*], the Pareto hazard increases faster than the Gompertz hazard (that is, with the Gompertz curve extended into the interval [*N, U*]). This is illustrated in Fig. 2 for females and males separately. The point *x*_*A*_ is the point at which the rate of increase of the Pareto hazard changes from being slower than that of the Gompertz hazard, to being faster. So we see a *late life mortality deceleration* in [*N, x_A_*) together with an *advanced age mortality acceleration* in (*x*_*A*_, *U*]. These are features consistent with a *finite upper bound* to the lifetimes distribution. This situation occurs in all the Netherlands and Australian cohorts. There is another point, *x*_*B*_, at which the hazards are equal, also given in Tables 2 and 3, and illustrated in Fig. 2. Other locations of interest, also given in Tables 2 and 3, are the point of inflection of the Gompertz, *x*_*I*_, and the point *x*_*C*_ where the first derivatives of the log hazards are equal for Gompertz and Pareto. Estimated parameters 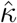 and *Û* along with the standard errors for the Australian cohorts are in Tables 4 and 5.

The female 1860 cohort has an outlier in *Û* due to some exceptionally low mortalities at extreme ages. Refitting the STLT model with these omitted gives *Û* = 110.7650 (*SE* = 0.653) in line with the other cohorts.

### The mixture model for lifetimes

To formulate this model we concentrate on the extreme right hand end of the mortality distribution, that is, for ages above the transition age *N*. Write the lifetime *T* of a person in the population as *T* = *N* + *Y*, where *Y* has the distribution of *T* conditional on *T > N*, thus, from Eq. (3), for given *κ, θ >* 0,

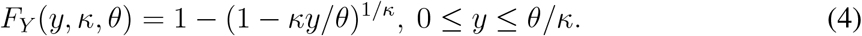

Assume now that associated with person *i* is a positive random variable *K*_*i*_ whose realized value is *κ* in Eq. (4); thus, *Y*_*i*_ for person *i* has the distribution in Eq. (4), conditional on the given value of *K*_*i*_:

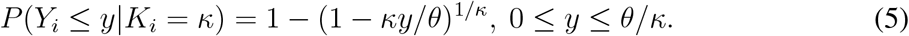

Assume a density *f*_*K*_(*κ*), *κ >* 0, for the *K*_*i*_ and integrate Eq. (5) with respect to it to get the marginal cdf for a typical member of the population as

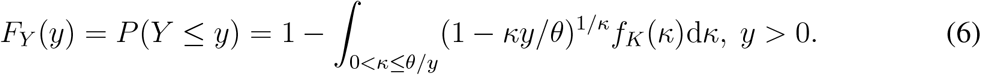

Differentiating Eq. (5) gives the conditional density of *Y* as

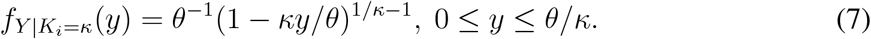

The conditional expectation calculated from Eq. (7) is *E*(*Y |K* = *κ*) = *θ*(*κ* + 1)^*−1*^, showing that, as *κ* increases, the expected lifetime decreases. Persons with higher values of *κ* have shorter lifetimes, on average. As *κ* approaches 0 lifetimes become longer and would be infinite for *κ* = 0.

Inspection of the Netherlands and Australian data in Fig. 5 suggests that the *κ* values might be approximately modeled as a lognormal or gamma distribution. We found a reasonably good fit of a gamma distribution. The limited number of observations on *κ* will increase as more data and analyses become available.

Suppose now in our setup that the *K*_*i*_ have a gamma distribution with parameters *a, b >* 0 and density

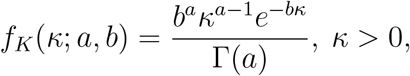

as shown in Fig. 5. Consequently, overall, the *Y*_*i*_ are taken to be independent and identically distributed with marginal density

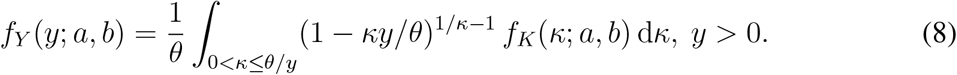

Parameters *θ* and *N* are specific to the cohort under consideration.

### Observed vs expected centenarians

Table 6 gives the observed and expected numbers in the cohorts living to ages 100+, 105+, and 110+, according to the mixture model.

**Table 6:** Observed (expected) number alive, by age and cohort. In the “All” row, the expected numbers were produced using the “All” estimates in Tables 2 and 3 for the calculation in Eq (1) in the main text. Approximate 95% confidence intervals around the estimated numbers were calculated by a resampling method.

| Cohort | Females |  |  | Males |  |  |
| --- | --- | --- | --- | --- | --- | --- |
|  | 100+ | 105+ | 110+ | 100+ | 105+ | 110+ |
| 1893 | 381 (384.8) | 18 (27.2) | 0 (0.6) | 123 (189.7) | 5 (23.7) | 0 (1.7) |
| 1894 | 428 (402.3) | 28 (25.3) | 1 (0.4) | 132 (126.2) | 5 (10.0) | 0 (0.4) |
| 1895 | 418 (431.3) | 31 (27.5) | 0 (0.4) | 108 (192.0) | 5 (23.3) | 0 (1.6) |
| 1896 | 436 (437.2) | 17 (27.5) | 1 (0.4) | 116 (114.1) | 5 (3.3) | 0 (0.0) |
| 1897 | 412 (472.5) | 22 (35.7) | 0 (0.8) | 103 (106.8) | 2 (3.3) | 0 (0.0) |
| 1898 | 502 (488.6) | 22 (31.1) | 1 (0.5) | 96 (103.3) | 3 (2.8) | 0 (0.0) |
| 1899 | 496 (498.2) | 31 (31.8) | 3 (0.5) | 104 (113.9) | 4 (5.2) | 0 (0.0) |
| 1900 | 482 (483.6) | 28 (29.0) | 1 (0.4) | 111 (106.2) | 4 (3.7) | 0 (0.0) |
| 1901 | 478 (533.8) | 20 (38.3) | 0 (0.8) | 108 (121.8) | 3 (6.8) | 0 (0.1) |
| 1902 | 529 (544.6) | 32 (32.1) | 0 (0.4) | 110 (109.2) | 3 (3.4) | 0 (0.0) |
| 1903 | 595 (554.6) | 35 (29.7) | 0 (0.3) | 103 (97.7) | 10 (3.0) | 1 (0.0) |
| 1904 | 581 (623.6) | 28 (48.6) | 1 (1.3) | 119 (140.9) | 6 (8.4) | 0 (0.2) |
| 1905 | 610 (594.3) | 44 (37.4) | 0 (0.7) | 112 (112.1) | 2 (3.5) | 0 (0.0) |
| 1906 | 639 (589.4) | 52 (28.9) | 2 (0.3) | 99 (120.2) | 6 (4.9) | 0 (0.0) |
| 1907 | 586 (627.8) | 38 (37.8) | 0 (0.6) | 128 (114.8) | 11 (3.2) | 0 (0.0) |
| 1908 | 657 (769.5) | 37 (71.3) | 0 (2.7) | 99 (107.0) | 9 (3.6) | 0 (0.0) |
| All | 8230 (8151.7) | 483 (481.0) | 10 (6.6) | 1771 (1722.4) | 83 (55) | 1 (0.4) |

### The regression model for centenarians

Fig. 6 shows the raw data and regression predictions on linear and log scales. The regression function is

**Figure 6:**
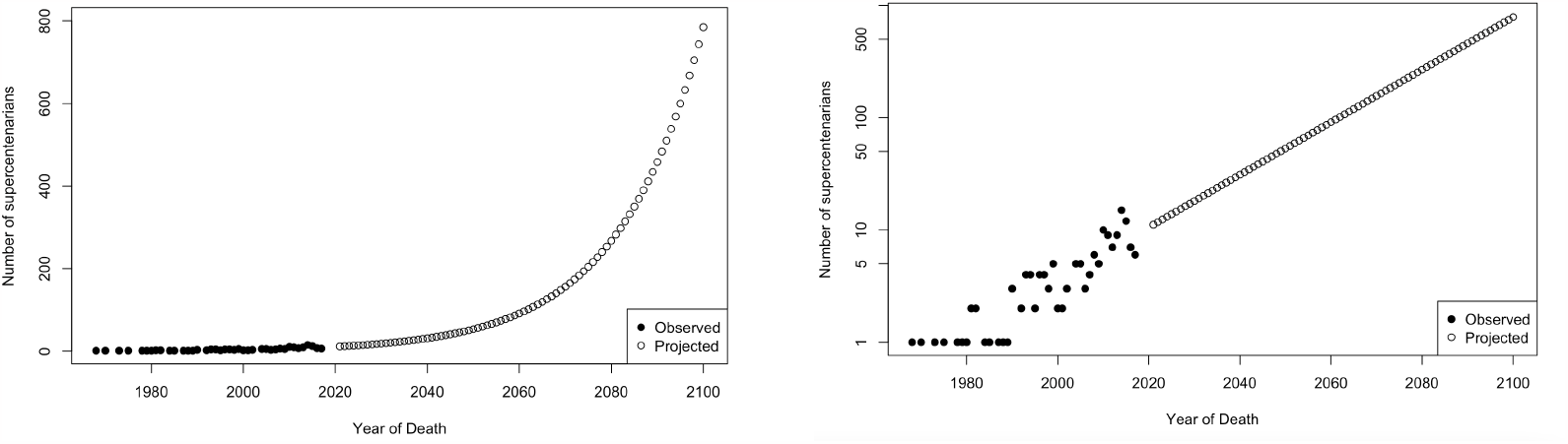
Regression models for centenarians. Left column, linear scale; Right column, log scale.

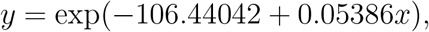

where *x* is calendar year above 2021 and *y* is the number of supercentenarians dying in that year.

## Data Availability

The data used in this paper are all publicly available. The Netherlands data is obtained from the Harvard Dataverse (*35*) and the Human Mortality Database (https://www.mortality.org/). More details about the data augmentation and quality checking can be found in (*5*). The Australian data is obtained from the Human Mortality Database (https://www.mortality.org/) (*6*). The England and Wales supercentenarians data is obtained from the International Database on Longevity (https://www.supercentenarians.org/).

## Code Availability

Code scripts used in this paper are available via the GitHub webpage:

https://github.com/feihuang921/hmmn.git.

The STLT model is implemented in R using the package published in GitHub and available at

https://github.com/u5838836/STLT.

